# Remodeling of ryanodine receptor isoform 1 channel regulates the sweet and umami perception of *Rattus norvegicus*

**DOI:** 10.1101/2021.07.20.453074

**Authors:** Wenli Wang, Dingqiang Lu, Qiuda Xu, Yulian Jin, Guangchang Gang, Yuan Liu

## Abstract

Sweet and umami are respectively elicited by sweet/umami receptor on the tongue and palate epithelium. However, the molecular machinery allowing to taste reaction remains incompletely understood. Through a phosphoproteomic approach, we found the key proteins that trigger taste mechanisms based on the phosphorylation cascades. Thereinto, ryanodine receptor isoform 1 (RYR1) was further verified by sensor and behaviors assay. A model proposing RYR1-mediated sweet/umami signaling: RYR1 channel which mediates Ca^2+^ release from the endoplasmic reticulum is closed by its dephosphorylation in the bud tissue after umami/sweet treatment. And the alteration of Ca^2+^ content in the cytosol induces a transient membrane depolarization and generates cell current for taste signaling transduction. We demonstrate that RYR1 is a new channel in regulation of sweet/umami signaling transduction and also propose a “metabolic clock” notion based on sweet/umami sensing. Our study provides a rich fundamental for a system-level understanding of taste perception mechanism.

## 1. Introduction

Accurate perception of taste information in food is crucial for human and other animal survival and evaluating food quality. Animals develop a strong preference for sugar and amino acid/peptide as the fundamental source of energy through TCA cycle and oxidative phosphorylation. Sweet/umami (sugar and amino acid/peptide) as reward taste can be elicited by their receptor cells on the tongue and palate epithelium [1]. To understand taste signal information, many studies focused on the sweet/umami receptors and the related signaling pathways by live cell imaging, genetics, bioinformatics and behavioral studies [1–3]. T1Rs, a family of G-protein-coupled receptors (GPCRs) as the mammalian taste receptors have been identified in sweet and umami detection [4–6]. Despite relying on different receptors, sweet and umami transduction use a common signaling pathway and converge on common signaling molecules, such as transient receptor potential cation channel subfamily M member (TRPM4 and TRPM5), type 3 inositol-1,4,5-trisphosphate receptors (IP3Rs) and phospholipase C (PLCβ2), calcium homeostasis modulator 1 (CALHM1) [3, 7–10]. CALHM1 is a voltage-gated ATP-release channel required for sweet and umami taste perception [10]. IP3Rs are the principal mediator of sweet and umami taste perception and would link phospholipase PLCβ2 to TRPM5 activation [9]. Moreover, sweet and umami perception in type II taste receptor cells on the tongue, both relies on the activity of TRPM4 and TRPM5, which is activated by the change of intracellular calcium (Ca^2+^) [3, 8, 11, 12]. Together, these studies demonstrated the similarity of sweet and umami in signaling pathway. However, the molecular machines allowing to taste reaction remain incompletely understood. Here, a fundamental question that we should address is whether new receptors/molecules activate TRPM4/TRPM5 and further mediate the signaling transduction of sweet and umami.

An intriguing, universal link exists between the Ca^2+^-mediated signaling and sweet, umami and bitter taste signaling. IP3Rs and ryanodine receptors (RyRs) are the two major calcium (Ca^2+^) release channels on the sarco/endoplasmic reticulum [13]. IP3Rs as the linkers of phospholipase PLCβ2 and TRPM5 have been demonstrated in sweet and umami taste perception [9]. Ryanodine receptor isoform 1 (RYR1) is a key mediator of the cellular regulation of calcium homeostasis, modulating multiple intracellular signaling pathways and physiological functions, most importantly in the sarcoplasmic reticulum required for skeletal muscle contract [14]. Ryanodine receptors have verified in the mammalian peripheral Type II and Type III taste receptor cells [15]. RYR1 was selectively associated with L type calcium channels in mammalian taste cells, and they can interact in neurons like in muscle[16]. However, how RYR1 mediates sweet and umami signal transduction remains poorly understood.

Protein phosphorylation is a mechanism of regulation that is extremely important in complex signal transduction network [17]. Phosphoproteomic as an essential approach is used to understand the complex molecular circuitry of cellular signal transmission [18, 19]. To make a thorough inquiry on the mechanism of taste (sweet/umami) perception, we performed a phosphoproteomic analysis for the rat bud tissue after 8 typical taste stimuli (sucrose, sucralose, aspartame, thaumatin, succinic acid disodium salt (SUC), monosodium glutamate (MSG), guanosine 5’-monophosphate disodium salt (GMP) and disodium lnosinate (IMP)) treatment (Figure. 1). Before phosphoproteomic analysis, these stimuli concentrations were measured based on taste bud tissue sensor. We discover a role for RYR1 channel in sweet and umami signal transduction, and this is a previously uncharacterized point that regulates taste perception. In addition to providing insights into sweet/umami signal transduction pathways, these data will serve as an important resource for further investigation of nutrient sensing mechanism.

**Fig. 1.**
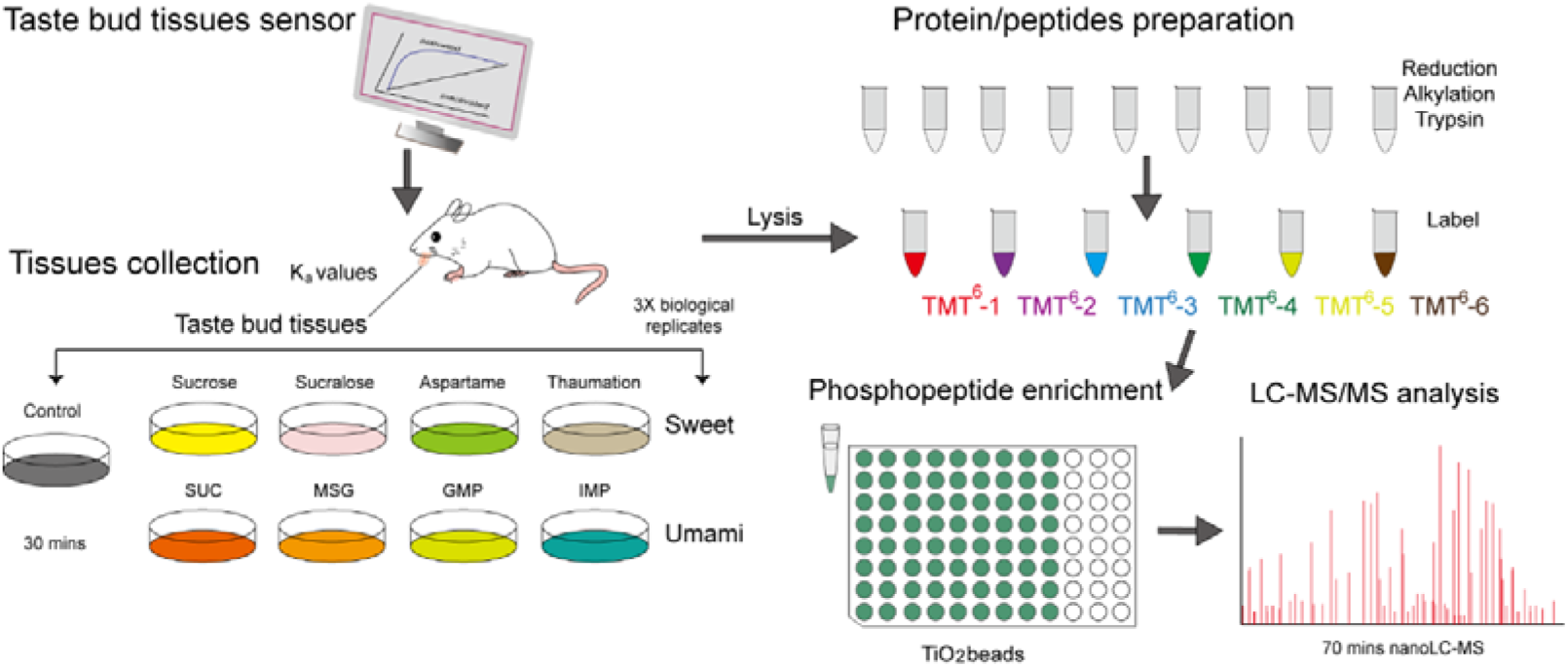
The schematic diagram of approach to protein phosphorylation analysis in rat bud tissue after sweet and umami stimuli

## 2. Methods

### 2.1 Material

The 8∼10 weeks old Sprague Dawley (SD) rats weighed about 200 g and were all male provided by Shanghai Jiesijie Laboratory Animal Co., Ltd. (Shanghai, China).

### 2.2 K_a_ values measurement of 8 taste stimuli based on taste bud tissue sensor

After the SD rats were sacrificed by decapitation, the rat taste bud tissue (from the tip to foliate papillae) was collected and fixed between two nuclear microporous membranes using the sodium alginate-starch gel as a fixing agent as previously described [20]. Due to the gelatinization of sodium alginate solution, this sandwich-type membrane was immersed into 5.0% CaCl_2_ solution for 10 s to form the sensing membrane and then fixed on the pretreated glassy carbon electrode. The biosensor with a three-electrode system (a glassy electrode fixed with a taste-measuring membrane as a working electrode, an Ag/AgCl electrode as a reference electrode, and a platinum-wire electrode as a counter electrode) provided a desired microenvironment for ligand-receptor binding.

The action potential generated by the taste bud cells after 8 stimuli treatment can be transmitted as electrical signals to the signal acquisition system. The amperometric *i-t* curve method was used to measure the response current of 8 stimuli at a voltage of 0.4 V, and ultrapure water as the base fluid was tested. The current change rate was calculated based on the steady state current values at the same time point before and after simulation. Due to substrate saturation effects, the linear concentration range and the kinetic curve of 8 taste compounds were determined, respectively. Their K_a_ values were calculated according to the method of Wei [21].

### 2.3 Tissues collection

All rats were done in compliance with the regulations and guidelines of Shanghai Ocean University institutional animal care and conducted according to the AAALAC and the IACUC guidelines. After jejunitas treatment 12 h, taste bud tissue (tongue) was quickly collected and respectively placed in 9% saline solution (including taste stimuli) for 30 min at room temperature, and 9% saline solution was as control in whole experiments. The concentrations of 8 stimuli were defined basing on their K_a_ values in this study. All samples after treatment were collected for further analysis.

### 2.4 Protein/peptide preparation

The tissues were cut into pieces and lysed in lysis buffer (100 mM NaCl, 50 mM Tris-HCl pH 7.4, 1% Triton X-100 (v/v) and 1 mM phenyl-methylsulfonyl fluoride). The samples were sonicated with an ultrasound for 100 cycles of 20 min at 4 °C, and then the supernatant was diluted 1:5 with ice-cold acetone and stored at −20 °C for overnight. The pellets were collected by centrifugation again for 10 min at 12,000 g at 4°C. The above two steps were repeated. The total pellets were dried and their protein concentrations were determined using a Bradford assay (Bio-Rad). The proteins were reduced and alkylated in the buffer containing 100 mM triethyl ammonium bicarbonate (TEAB) and 10 mM Tris (2-carboxyethyl) phosphine (TECP) and 20 mM iodoacetamide. Proteins were digested overnight at 37 °C with trypsin (Promega) with an enzyme: substrate ratio of 1:40. and trypsin hydrolysates were labeled using tandem mass tags (TMT) reagent in acetonitrile for 1 h at 25 °C. Samples were dried down and subjected to phosphopeptides analysis.

### 2.5 Phosphopeptides enrichment and reversed phase nLC-MS/MS analysis

Phosphorylated peptides were enriched using phosphopeptides buffer according to the manufacturer’s protocol (TiO_2_ beads, GL Sciences, Japan). Phosphorylated peptides were dried down and dissolved in 200 μL of Nano-RPLC buffer A consisting of 2% ACN/0.1% FA. These peptides were subjected to reversed phase LC-MS/MS analysis using Easy-nLC 1000 system coupled to an Orbitrap Q Exactive Plus mass spectrometer (both Thermo Scientific) for the phosphoproteome analysis. The Nano-RPLC system was equipped with samples a trapping column (Thermofisher Dionex, PepMap100, C18, 3 μm, 3 cm ×100 μm) and an analytical column (Thermofisher Dionex, PepMap100, C18, 3 μm, 15 cm ×75 μm). Trapping was performed for 10 min in solvent A at 2 μL/min. An elution gradient of 5-35% acetonitrile (0.1% formic acid) for 70 mins was used for the phosphoproteome samples analysis. The mass spectrometer was operated via Q Exactive (Thermo Scientific) in data-dependent mode. The Orbitrap Q Exactive Plus full-scan MS spectra from m/z 300–1800 were acquired at a resolution of 70000 for MS and 17500 for MS/MS. Up to 20 most intense precursor ions were selected for fragmentation. HCD fragmentation was performed at normalised collision energy of 25%.

### 2.6 Data analysis of phosphoproteomics

Raw files were processed using Proteome Discoverer™ (version 1.3) using the SEQUEST® search engine. The data search was performed against the complete rat Swiss-Prot database. Default settings were used, with the following minor changes: methionine oxidation, protein N-term acetylation, protein N-term pyro-glutamate, deamidation of asparagine and glutamine, and methylation of cysteine as variable modifications. TMT isobaric labeling (+229.163 Da) of lysine in protein N-term were set as static modifications. A false discovery rate (FDR) of 1% was applied at the protein, peptide, and modification level.

Bioinformatics analysis was performed with Origin 9.1, Microsoft Excel and R statistical computing software. Annotations were downloaded and extracted from UniProtKB, Gene Ontology (GO), and Kyoto Encyclopedia of Genes and Genomes (KEGG). In addition, protein-protein interaction networks were obtained by STRING database (http://string.embl.de/). All figures were finally organized in Adobe Illustrator CS6.

### 2.7 The role of RYR1 based on taste bud tissue sensor

According to the previous preparation method of sensor[20, 22], the rat taste bud tissue from SD rat was collected. Using the sodium alginate-starch gel as a fixing agent, taste bud tissues were fixed between two nuclear microporous membranes, and they were made into a sandwich structure. This prepared membrane structure was immersed in 5 g/100 mL CaCl_2_ solution for 10 s to fix them well. Then physiological saline was used to remove remaining Cl^−^, Ca^2+^ on the membrane. The fixed membrane structure containing taste bud tissue was respectively placed in the ryanodine solution with different concentrations (10^−5^∼10^−6^ mol/L) for 30 min and then fixed on the surface of glassy carbon electrode mentioned above. This sensor was determined by amperometric *i-t* curve method in a three-electrode system, and then the response current was plotted for a series of MSG solution (10^−6^∼10^−12^ mol/L). The rate of current change as detection index was used to calculate the role of RYR1 in the response of MSG to taste bud tissue.

### 2.8 Taste behavioral assays

Taste behavior was assayed using a short-term assay that directly measures taste preferences by counting water intake volume. On testing days, each three rats were placed into the cage, and presented with water and two different tastant solutions (300 mmol/L sucrose and 50 mmol/L MSG+1mmol/L IMP) [23]. Each 500 μL 1×10^−5^ mol/L ryanodine solution was smeared in the tongue of rat and 2× per day. The rats without ryanodine solution as the control groups were investigated in comparison to the treated groups. Water consumption was measured in each over 24 h. After an initial recording period of 24 h, the samples were renewed and the locations were switched to control for side bias. Water consumption was then measured over another 24 h. Tubes were of identical construction and all individual drinking tubes were tested over night before the experiment to ensure that they did not lose water through dripping.

### 2.9 Immunostaining of RYR1 in taste bud tissue

Rat taste bud tissues were fixed with a 4% paraformaldehyde solution. Paraffin-embedded tissues with 3 µm thick sections were deparaffinized with xylene for 3×30 minutes and incubated in graded concentration of ethanol and then washed with PBS for 3×5 minutes. The dewaxed and rehydrated sections were stained in hematoxylin solution until the nuclei becoming blue. The sections were washed twice with 70% ethanol, absolute ethanol and xylene for 10 min, respectively. The dewaxed and rehydrated samples of paraffin section were incubated in boiling 1 mM EDTA buffer (pH=8.0) for 2 min, and sections were washed with PBS with 0.1% Triton X-100 (PBST), blocked with 3% goat serum (Invitrogen, US) in PBST. Sections were incubated with primary antibody (rabbit anti-RYR1 antibody 1:500, AB9078, millipore) overnight at 4 °C, incubated with fluorescence-tagged secondary antibody (goat anti-rabbit IgG 1:1000, AB6721, USA) for 30 min at room temperature. The sections were examined using an Olympus light microscope (CX31, Olympus).

### 2.10 Double-label immunofluorescence for RYR1 and calnexin in bud tissue

The fixed tissues were placed in 30% sucrose solution overnight at 4 c for cryoprotection. Tissues were embedded in OCT compound and sectioned at 5 µm thickness on a cryostat. Sections were washed in PBS with 0.1% Triton X-100 (PBST), blocked with 10% donkey serum in PBST, incubated with primary antibodies (RYR1, 1:400, abcam2868, US and calnexin, 1:500, abcam22595, US) overnight at 4 °C, and incubated with the Alexa Fluor 488-conjugated donkey anti-mouse IgG (1:1000, abcam 150105, US) and Alexa Fluor 594-conjugated donkey anti-rabbit IgG (1:1000, abcam 150076, US) as the secondary antibodies for the RYR1 and calnexin antibodies, respectively. The sections were observed under a confocal laser scanning microscope (FV1200, Olympus, Japan). The Green fluorescence was excited with 488 nm and imaged through a 525nm long-path filter. The Red fluorescence was excited at 543 nm and imaged through a 590 nm long-path filter.

### 2.11 Statistical analysis

To ensure the consistency and reproducibility of our results, we conducted at least three repeated trials performing in different preparations and from at least three different animals.

## 3. Results

### 3.1 The allosteric constant of 8 taste stimuli for rat bud tissue

In consideration of the effect of taste stimuli concentration on protein phosphorylation, we measured the response ranges of diverse sweet and umami substances including protein, nucleotide, amino acids and sugar by rat bud tissue sensor. This sensor maintained the activity and integrality of taste bud cells so that the information of ligand-taste receptor binding can be well collected by electrical signals. The allosteric constants (K_a_) of ligand-receptor, which were defined as the concentration of ligand at half of the maximum output signals of cells, were calculated by the signal output generated by the interaction of the receptors in bud tissue to ligands (sweet or umami molecules)[20]. We found that this sensor provided the response signals for 8 taste stimuli at low concentrations (10^−13^ mol/L). The K_a_ values of 8 taste stimuli were listed in Supplementary table 1. These results implied that sweet/umami receptors in rat bud tissue had the excellent recognition and sensing effect to 8 taste stimuli.

The perception ability of four umami substances in rat taste bud tissue is: GMP>IMP>SUC>MSG. However, the sensing ability of umami hT1R1 towards the abovementioned four umami ligands was much lower than bud tissue[24]. This may be related to umami taste transduction mechanism based on multiple umami receptor (heterodimers T1R1/T1R3 and metabotropic glutamate receptors)[7] or the cross-talk among various signal pathways in bud cell[8, 11]. Artificial sweeteners can activate the same sweet taste receptor on the tongue, e.g., aspartame as artificial sweetener showed a certain sensing performance of sweet taste receptor nanovesicles-based bioelectronics tongues[25]. In this study, the perception ability of four sweet stimuli is: Sucrose>Aspartame>Sucralose>Thaumatin (Supplementary table 1). Together, to gain insight into sweet and umami taste signal networks whose activity is regulated by protein phosphorylation, we used the K_a_ values of 8 taste stimuli to respectively treat rat bud tissues for 30 min and collected these tissues for phosphoproteomic analysis.

### 3.2 Analysis of the differential phosphoproteins of stimulated taste bud tissue

To identify the relevant effectors involved in sweet and umami signal transduction, we used a proteomic approach to identify phosphoproteins and their phosphorylation sites in the stimulated taste bud tissue. Phosphoproteins were analyzed and identified by liquid chromatography tandem mass spectrometry (LC-MS/MS) combined with a *Rattus norvegicus* Uniprot database. Protein and peptide false discovery rates were set at 1%. We defined the identified phosphoproteins as differential proteins if there was a fold change in excess of 1.2 or in less of 0.83 proteins changed significantly in the stimulated taste bud tissues compared with control. For four sweet stimuli, we identified the total 196 differential proteins, which clearly revealed that substantial changes in the phosphorylation levels (Figure. 2a). Of these, 125 phosphoproteins (78 upregulated and 47 downregulated) were identified in the bud tissue of thaumatin treatment, containing 470 phosphorylation sites. More than 60% phosphoproteins contained more than one phosphorylation site, with some proteins having as many as 27 sites. Among other sweet molecules, aspartame showed 98 phosphoproteins (53 upregulated and 45 downregulated), whereas data from of sucrose and sucralose showed 61 and 38 phosphoproteins, respectively. 19 differential phosphoproteins as the shared ones were accounted in the bud tissues from four sweet stimuli by Venn analysis (Figure. 2b). Phosphorylation in the shared proteins was biased toward Ser compared Thr (76%: 17%), a similar with the findings in eukaryotes, where Ser phosphorylation may account for 80-90% of total phosphorylation sites[26].

**Fig. 2.**
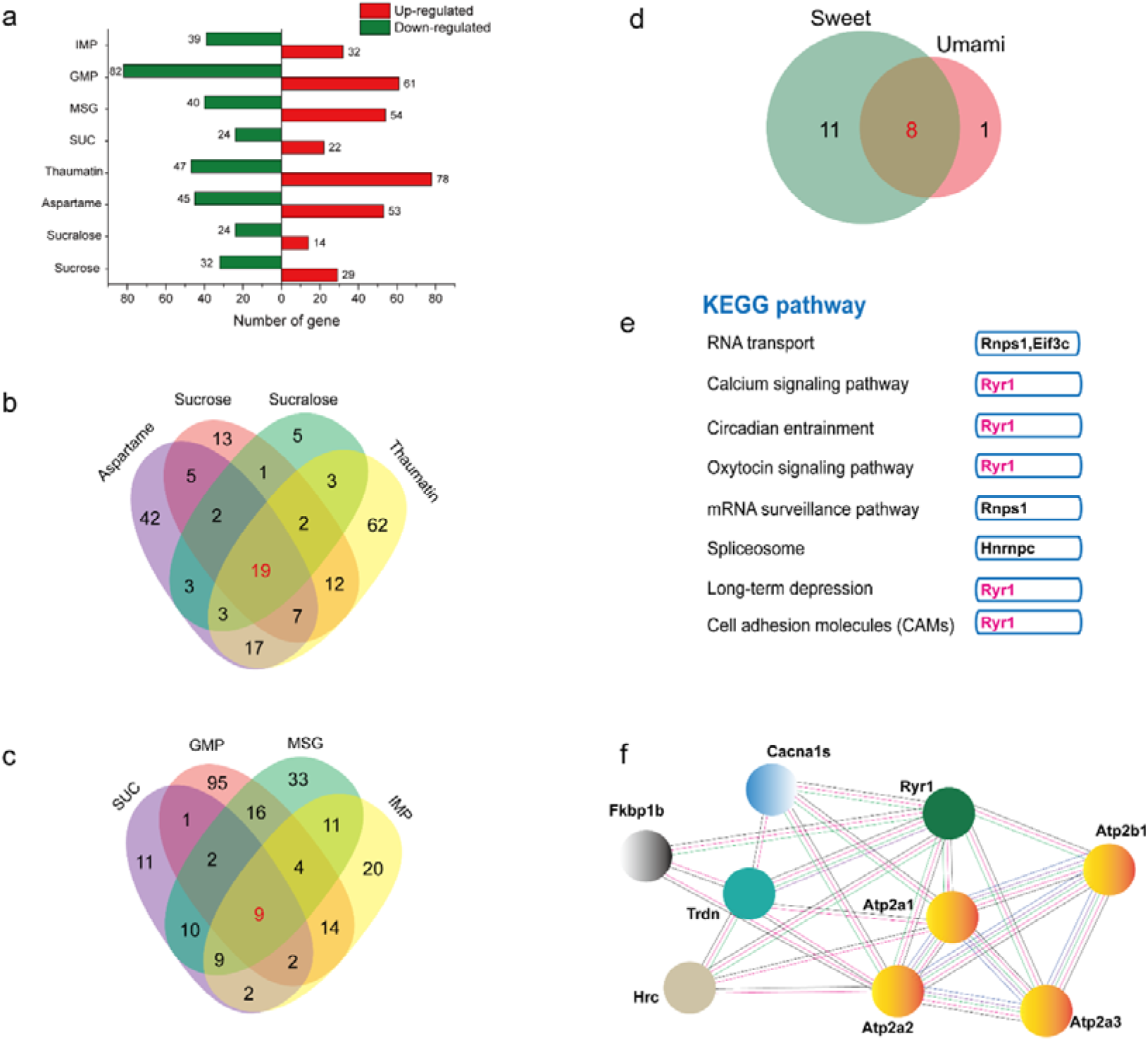
a: The information of differential genes in the rat bud tissues after 8 taste stimuli treatment including sucrose, sucralose, aspartame, thaumatin, succinic acid disodium salt (SUC), monosodium glutamate (MSG), guanosine 5’-monophosphate disodium salt (GMP) and disodium lnosinate (IMP); b: Venn diagram showing the distribution of sweet-related phosphoproteins in rat bud tissues; c: Venn diagram showing the distribution of umami-related phosphoproteins in rat bud tissues; d: Venn diagram showing 8 common differential proteins in all treated bud tissues; d: Top enriched pathways in 8 common proteins in the treated bud tissues; f: The related protein to RYR1 in taste bud tissue association predicted network by STRING (http://string-db.org).

Though there were many differential phosphoproteins in the bud tissue of thaumatin and aspartame stimuli, we still identified more differential phosphoproteins in the bud tissues from umami stimuli than sweet molecules. A total of 239 differential phosphoproteins were identified in the bud tissues from four umami stimuli compared with the unstimulated bud tissues (Figure. 2a). Of these, 143 differential phosphoproteins (61 upregulated and 82 downregulated) in the bud tissue of GMP treatment account for 60% of total differential phosphoproteins, containing 551 phosphorylation sites. Next, we examined 94 and 71 differential proteins in the bud tissue of MSG and IMP treatment, respectively. In contrast, the minimal amounts of differential phosphoproteins were identified in the bud tissue of SUC treatment. Closer inspection of these data, we found that protein phosphorylation of taste bud tissue would be easier in present of taste stimuli including amino-group. These data suggested an interesting insight regarding the 9 shared protein from four umami stimuli, and 8 downregulated phosphoproteins also were found in the bud tissues from four sweet stimuli as the shared ones (Figure. 2c&d).

Based on the studies of the Ca^2+^-mediated sweet (umami) taste signaling[5, 6, 27], the related differential phosphoproteins to Ca^2+^ channel should be summarized to elucidate sweet and umami signaling transduction (Table S4). The 8 taste stimuli were shown by a number of specific related phosphoproteins in the corresponding protein levels (Figure. 3). Several key proteins in Ca^2+^-mediated signaling were found in those taste bud tissues after sweet and umami stimulation, such as histidine-rich calcium binding protein (HRC), RYR1 and ATPases (Figure. 2e). It is particularly noteworthy that, RYR1 as a shared differential phosphoprotein was found in all the treated taste bud tissues.

**Fig. 3.**
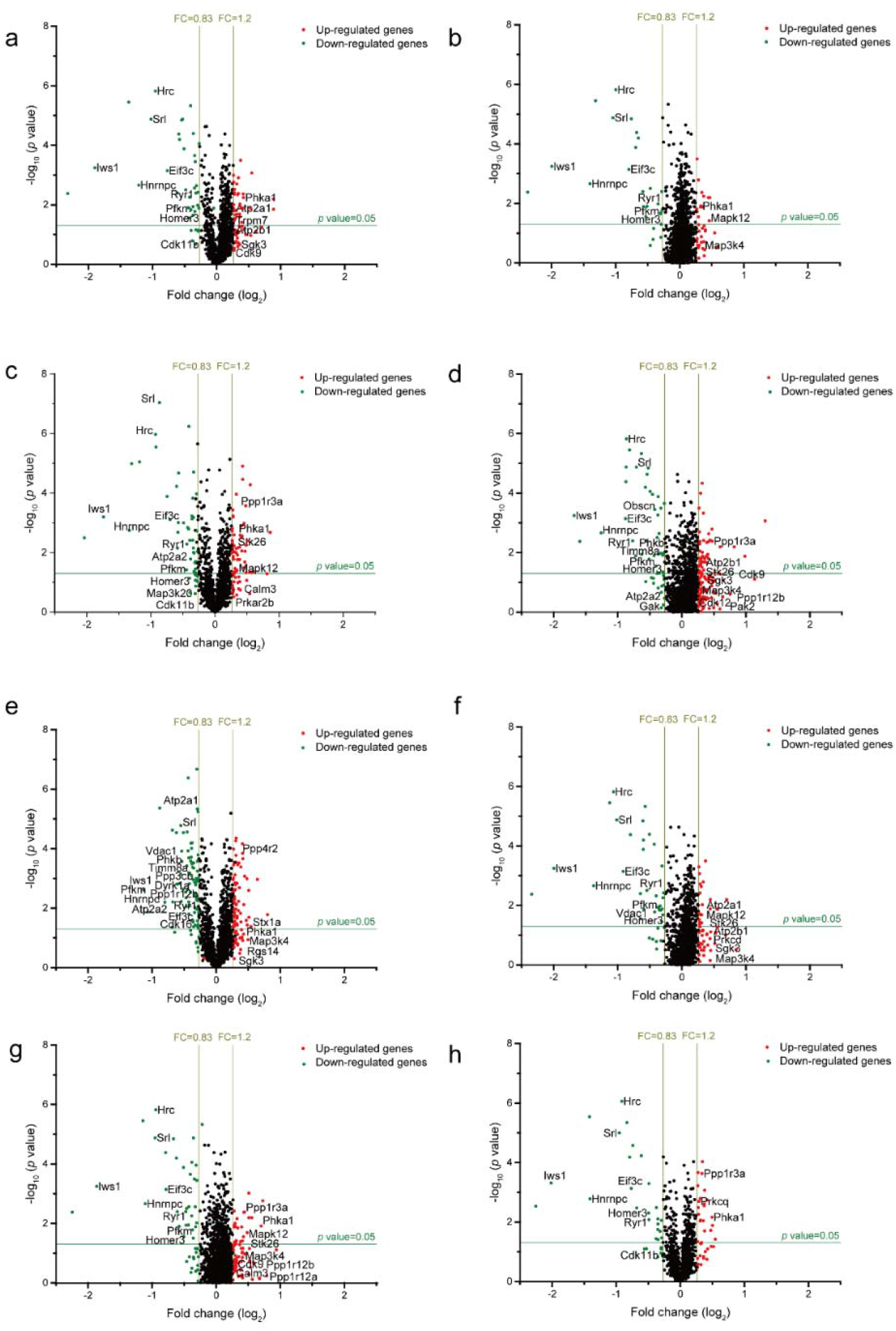
Volcano plots of protein samples from rat bud tissue treated with various sweet and umami substances. Compared to black sample, Fold changes in proteins abundance (Proteins passing both cutoffs) for each treated sample are shown as a function of statistical significance. Significance cutoffs were p-value□=□0.05, FC□=□1.2 and FC□=□0.83. Each protein is represented with a single data point, corresponding to the peptide with the lowest p-value. The common interactor of the bud tissues treated with 8 taste substances were highlighted in cyan and labeled with their gene names. a: Sucrose; b: Sucralose; c: Aspartame; d: Thaumatin; e: GMP; f: IMP; g: MSG; h: WSA

### 3.3 A series of key protein related to Ca^2+^-mediated sweet (umami) taste signaling

As mentioned above, these key proteins that may mediate the connectivity of umami/sweet signaling transductions, were further analyzed according to KEGG pathway analysis and STRING database (http://string.embl.de/) (Figure. 2e&f). Closer inspection of these data, however, provides interesting insights regarding the regulation of sweet and umami taste pathway by calcium signaling. RYR1, which mediates the release of from intracellular stores into the cytosol[14], is a shared differential phosphoprotein in all bud tissues after sweet and umami stimuli. HRC that can directly interact with sarco/endoplasmic reticulum Ca^2+^-ATPase (SERCA) to regulate Ca^2+^ homeostasis including the regulation of both Ca^2+^-uptake and Ca^2+^-release[28], also found in the mostly bud tissues after sweet and umami stimuli. Homer proteins regulate a number of Ca^2+^ handling proteins, including transient receptor potential channels and other Ca^2+^ permeable channels, ionotropic and metabotropic glutamate receptor, shank scaffolding proteins or endoplasmic reticulum (ER) Ca^2+^ channels[29]. Homer3 as an important differential phosphoprotein were found mostly (6/8) bud tissues samples after 8 taste stimuli treatment. Srl involved store-operated calcium entry (Gene Ontology: 0002115), is also a shared differential phosphoprotein.

In the perception of sweet and umami, despite relying on different receptors, their transductions converge on common signaling molecules[8], among which calcium homeostasis modulator 1 is expressed specifically in sweet/umami-sensing type II taste bud cell [10]. Thaumatin as a sweet protein activated calcium-sensing receptor with an EC_50_ value of 71 µM [30]. The previous studies demonstrated the similarity of sweet and umami in perception process which relies on the activity of TRPM5 and the relevant Ca^2+^signaling molecules, such as type 3 inositol-1,4,5-trisphosphate receptor, phospholipase C and calcium homeostasis modulator 1 [7–10]. The shared proteins RYR1 as an intracellular Ca^2+^ release channel has a functional coupling with IP3R in regulating intracellular Ca^2+^ signaling in skeletal muscle[31]. Nevertheless, IP3R is a common signaling molecule in sweet and umami transduction signaling pathway[9]. In current study, a series of key protein related to Ca^2+^-mediated signaling demonstrated a critical role of RYR1 in controlling sweet (umami) taste perception.

### 3.4 The roles of RYR1 in MSG perception based on rat taste bud tissue sensor

We reasoned that RYR1 as a key channel regulates Ca^2+^ signaling after sweet (umami) stimuli. Based on previous study, electrochemical signal generated by the interconnect allosteric cascade between the receptor and its ligand can truly reflect the biological effects of cell signaling amplification and transmission[20, 22, 24]. To prove the role of RYR1 in sweet (umami) perception, we further assembled a series of rat taste bud tissues with ryanodine at the concentrations of 10^−5^∼10^−6^ mol/L and constructed their tissue sensors. The changing rate of response current showed a decreasing relationship with the concentration of ryanodine comparison to the biosensor without assembled ryanodine (Figure. 4a). Increasing the concentration of ryanodine tended to decrease the response strength of MSG on sensor. As predicted, the ryanodine as an inhibitor of rat taste bud tissue has been interpreted when the concentration of MSG was at 10^−12^∼10^−6^ mol/L. The response of MSG to taste bud tissue was fully reduced in the presence of 4×10^−6^ mol/L ryanodine. Our result indicated that ryanodine inhibited the activity of RYR1 and further slowed down umami signaling. This finding suggested that RYR1 was required to keep umami signaling pathway integrality.

**Fig. 4.**
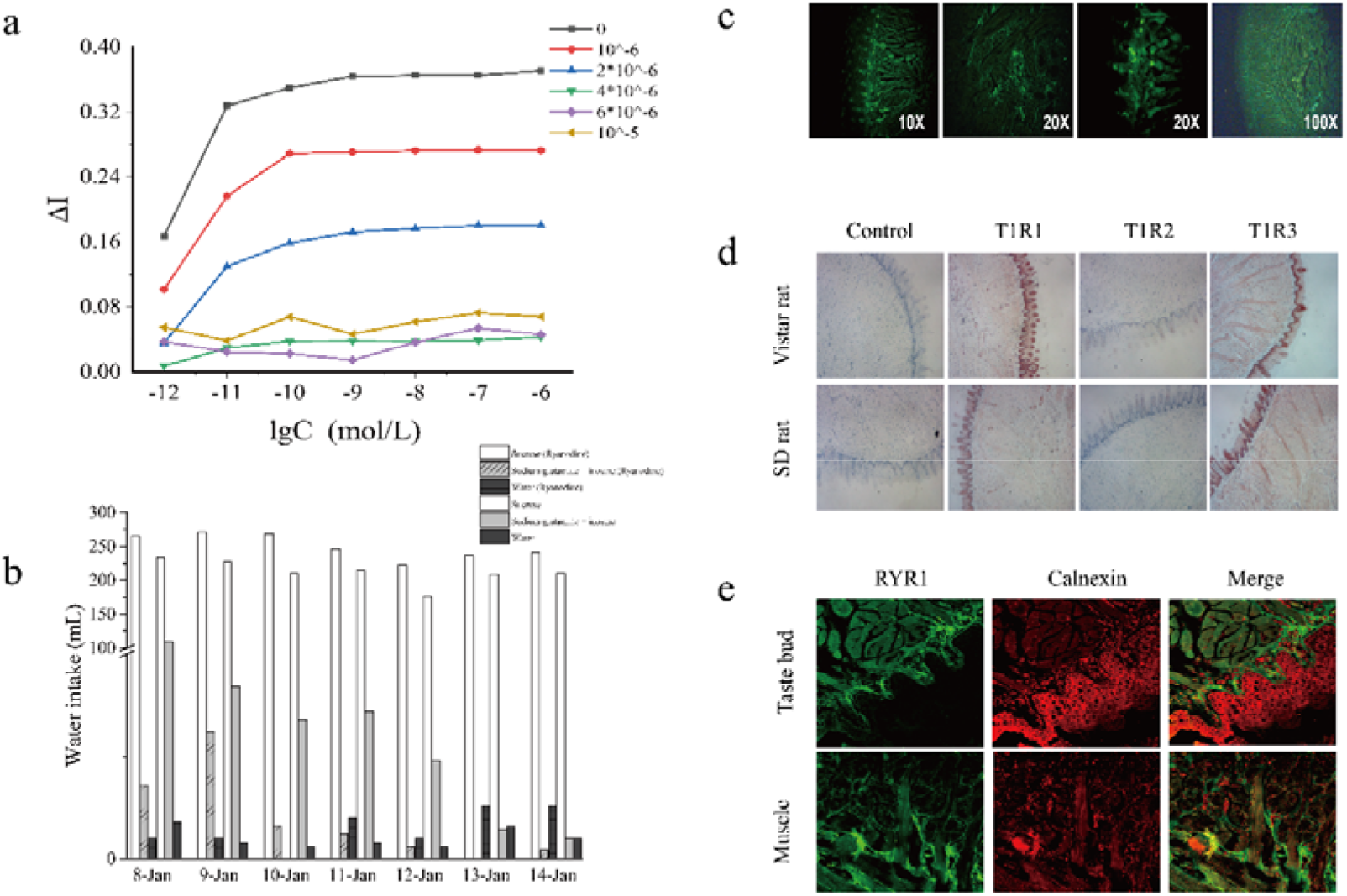
The confirmation of RYR1 roles in sweet and umami perception. a: The response of the taste bud tissue after ryanodine treatment to MSG solution (10^−12^∼10^−6^ mol/L) based on rat taste bud tissue sensor; b: The influence of RYR1 on taste perception based on behavioral tests (water intake volume); c: Expression profiling of fluorescently labelled RYR1 protein; d: Expression profiling of sweet and umami receptor (T1R1, T1R2 and T1R3) in two rat taste bud tissue; e: Confocal microscopy images of immunostaining illustrating the expression of RYR1 (green) and calnexin (red) in bud tissue and skeletal muscle.

### 3.5 The effective of RYR1 on behavioral tests

To further investigate the influence of RYR1 on taste perception, rats were exposed to different chemical stimuli (sucrose, IMP and MSG) to monitor their behavior. Rats, which was smeared in the tongue with 1×10^−5^ mol/L ryanodine solution, were presented with water, 300 mM sucrose and 50 mM MSG+1 mM IMP, the unsmeared rats as wild-type control group. Consistent with the sensor results, we examined the water intake volume on umami solution for the rats after ryanodine treatment has a decreasing trend than control ones (Figure. 4b). However, there is an interesting result that the water intake volume on sucrose solution for the rats after ryanodine treatment showed an increasing. In order to explain this reason and avoid the effect of chemical concentration on the water intake volume, we performed the experiment at a lower concentration (sucrose 1×10^−7^ mol/L, IMP 1×10^−8^ mol/L+ MSG 5×10^−7^) based on sensory pretest. The simulation results showed the total water intake volume for the rats after ryanodine treatment has a decreasing trend than control ones. Our results showed that sensitivity (the water intake volume) to sweet and umami was markedly inhibited in rats with ryanodine treatment. The results described above validated RYR1 as a key mediator is important in influencing taste perception for both sweet and umami.

### 3.6 Expression profiling of RYR1 and the related protein in the tongue

Ryanodine receptors (RyRs) with extraordinary size and complexity locate in the sarco/endoplasmic reticula (SR/ER) of most metazoan cell types[14]. The previous studies showed RYR1 required for skeletal muscle contraction is the predominant isoform[14]. Though calcium homeostasis modulators (CALHMs) are abundantly expressed in taste bud cells and have an important role in sensing sweet, bitter and umami[10]. Considering the result of RYR1 in the present manuscript, it is likely needed to observe the distribution of RYR1 in rat taste bud tissue. The immunohistochemical assay confirmed robust activation of RYR1 in rat bud tissue. (Figure. 4c), suggesting a mechanism for RYR1 channel gating by Ca^2+^, is essential for numerous cellular functions including sweet/umami signaling transduction in taste bud tissue.

Sweet and umami substances, which both provide sugar and amino acid/peptides, are the fundamental sources of energy for animals. Several studies have implicated a role for calnexin in controlling mitochondrial metabolism by Ca^2+^ flux, thereby regulating the energetic output of the oxidative phosphorylation pathway [32, 33]. To further investigate a potential role of calnexin for RYR1 mediated taste signaling, we also confirmed a predominant co-location of RYR1 with calnexin in the rat bud tissue and the tongue muscle via confocal microscopy (Figure. 4e). An example is the abundant calnexin, which stimulated the ATPase activity of sarco/endoplasmic reticulum Ca^2+^ ATPase (SERCA) [32]. Combined with the results of phosphoproteome, the two proteins most significantly associated with ATPase activity of SERCA. This finding suggested that calnexin was required in taste bud tissue to keep the activity of SERCA and regulate voltage-gated Ca^2+^ channels for sweet and umami perception. These findings together strongly support a notion that RYR1 related to Ca^2+^ signaling transduction proteins play a role in controlling Ca^2+^ flux in taste bud tissue following sweet and umami-tasting compound ingestion.

## 4. Discussion

Based on multiple signaling pathways, potential taste chemicals are precepted by interacting with taste receptor cells. Two distinct salt transduction mechanisms which are the epithelial sodium channel ENaC identified as the underlying receptor for mediating low-salt attraction and the amiloride-insensitive high salt transduction in Type II TRCs dependent on the presence of the Cl^−^ anion, have been reported [34, 35]. Recent study showed that a subset of excitatory neurons in the pre-locus coeruleus express prodynorphin, and these neurons are a critical neural substrate for sodium-intake behavior [36]. Acids are detected by type III taste receptor cells (TRCs), and the Otop1 expressed in type III TRCs has been proposed as a candidate sour receptor[37]. Bitter, sweet and umami stimuli are usually detected by the type II cells which express taste receptor type 1 (T1R) [5, 6] or T2R [38]. In addition, broadly responsive taste cell, as a subset of type III cells respond to sour stimuli but also use a PLCβ signaling pathway to respond to bitter, sweet and umami stimuli [39]. Calcium homeostasis modulator 1 (CALHM1) is expressed specifically in type II taste bud cells, which is a voltage-gated ATP-release channel required for sweet, umami and bitter taste perception [10]. Their transductions converge on the common signaling proteins (likes PLCβ2, IP_3_ and TRPM4/TRPM5 selectively expressed in taste tissue), despite relying on different receptors [8].

Sweet (sugar) and umami (amino acid/peptides) compounds provide energy (ATP, GTP and carbon) for animals via oxidative phosphorylation, TCA cycle and glycolysis. Sweet and umami stimuli are detected in the same cells [40] and share a common subunit T1R3 [4]. In spite of their diversity, T1Rs converge on a common intracellular signaling pathway, and their transporters couple to Ca^2+^ signaling. Sweet and umami stimuli interact with receptors to generate second messengers, or taste stimuli itself is transported into the cytoplasm of taste bud cell and activates downstream events[41]. When T1Rs are activated by sweet or umami stimuli, Gβγ dimers (belong to the family of GPCRs) are released, which stimulates PLCβ2 to mobilize intracellular Ca^2+^. The elevated Ca^2+^ levels in the cytosol lead to the opening of TRPM5 that effectively depolarizes taste cell. In this study, the key differential protein and their possible roles in taste are discussed below.

We found a shared phosphoprotein RYR1, which is a well-known family of ligand-gated channels responsible for the release of Ca^2+^ from the intracellular Ca^2+^ stores. A number of proteins-calmodulin, homer proteins and RYR1 stabilizing subunit calstabin1 (FKBP12)-bind and regulate RYR1 open-closed structure [42–44]. The binding of FKBP12 to RYR1 promoted and stabilized the closed state of the RYR1 calcium channel required for excitation-contraction coupling in skeletal muscle, but FKBP12-depleted, PKA-hyperphosphorylated, S-nitrosylation of RYR1 resulted in Ca^2+^ leak via the opening of RYR1 channels[44]. In this study, RYR1 containing S2840 and T4464 sites is progressively dephosphorylated in all treated taste bud tissues (Table. S3). Moreover, the RYR1 related proteins show a significant down change, with the downregulation of several kinases and the upregulation of phosphatase (Table. S4). We infer that the RYR1 channel exhibited weak “leaky” channel behavior (open state) when animals get hungry; during this time, Ca^2+^ slowly releases into the cytosol via RYR1 channel. However, the RYR1 channel in bud tissue would be closed after sweet and umami substances ingestion (Figure. 5). The alteration of Ca^2+^ content in the cytosol can induce a transient membrane depolarization. TRPM5, which responds to the rate of change in Ca^2+^ concentration, is directly activated at a certain range to generate significant cell current for taste signal transduction[12].

**Fig. 5.**
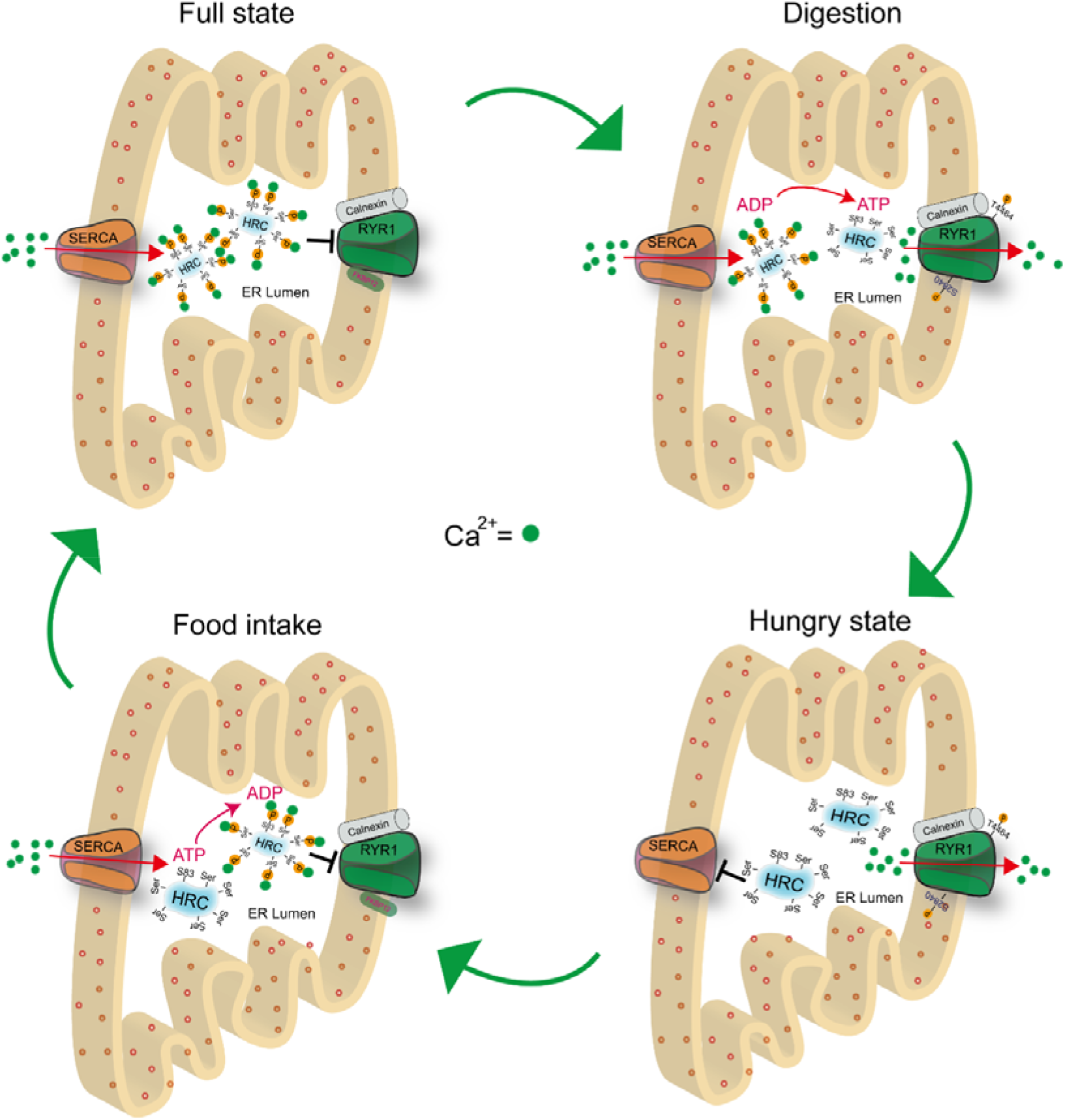
Model proposing the phosphorylation of HRC and RYR1-mediated sweet and umami signaling in metabolic cycles. After sweet and umami substances intake (Full state), RYR1 channel is closed in virtue of its dephosphorylation, and opening SERCA pump is regulated by HRC-p to increase Ca^2+^ flux from the cytosol to the ER. The digestion period of nutrient, RYR1 channel is opened by its phosphorylation, accompanied by Ca^2+^ flux from the ER to the cytosol and the dephosphorylation of HRC. The body goes hungry state until HRC in the ER was fully dephosphorylated, and SERCA pump is closed. SERCA pump is opened again by HRC-p to increase Ca^2+^ flux from the cytosol to the ER, and RYR1 channel is closed again since food intake beginning.

Histidine-rich Ca-binding protein (HRC) as the key differential protein also was found in the present study. HRC as the regulator of SERCA2a and RYR2 mediates SR Ca^2+^ homeostasis[28]. Moreover, the phosphorylation of HRC on Ser96 by Fam20C regulated the interactions of HRC with SERCA2a, suggesting a new mechanism for regulation of SR Ca^2+^ homeostasis[45]. In this study, we found the HRC containing 25 phosphorylation sites was robustly dephosphorylated, suggesting its role in regulating Ca^2+^ cycling that is critical for sweet and umami signaling. The above clusters also recruit other proteins implicated in the regulation of Ca^2+^ signaling, for example, the sarcoendoplasmic reticulum Ca^2+^-ATPase (SERCA) is identified as a differential protein. The activity of SERCA was stimulated by chaperone calnexin, which increased the ER Ca^2+^ content and controls mitochondrial Ca^2+^ responses[32]. Thus, the calnexin in rat bud tissue probably control the flux of Ca^2+^ in taste signaling by regulating the activity of SERCA. In addition, the shared differential protein sarcalumenin (SAR), a Ca^2+^-binding glycoprotein located in the longitudinal SR, regulates Ca^2+^ reuptake by interacting with SERCA[46]. The study has demonstrated SAR is essential for maintaining the Ca^2+^ transport activity of SERCA2a[46].

Changes in Ca^2+^ content in the cytosol via the regulation of RYR1 may due to the related protein amounts or activities. Homers are scaffolding proteins that bind GPCRs, IP3 receptors, RYR1, PLCβ and TRP channels through the CC region and selection of specific polyproline-rich sequences (PPXF) [29, 47]. Homer that cross-link G protein-coupled metabotropic glutamate receptor (mGluRs) is enriched at the postsynaptic density [48]. Homer3 was phosphorylated by CaMKII and activated by glutamate stimuli, resulting in attenuation of the physical and functional coupling between mGluR1 and IP3R [49]. Moreover, activity-induced Homer proteins further dynamically regulate the communication between calcium signaling related proteins to achieve the facilitation of calcium currents in neurons[50]. Homer3 as a key differential protein also was found based on the result of phosphorproteome, so its phosphorylation state regulates the physical coupling between mGluR1 and IP3R1 which partly reflects the Ca^2+^ signal during sweet and umami signaling pathway. Our data do not exclude the possibility of other Ca^2+^-dependent pathway by the induction by sweet and umami stimuli.

## 5. Conclusion

In summary, we have identified the key proteins in the rat bud tissue after sweet and umami stimuli by a phosphoproteomic approach. Our findings reveal a molecular mechanism in sweet and umami perception by a serial of RYR1 related phosphoproteins. For the taste bud tissue treated by sweet and umami substances, SERCA pump is activated by the regulation of dephosphorylated HRC and SAR to increase Ca^2+^ flux from the cytosol to the ER, and the close of RYR1 channel by dephosphorylation to stop the ER Ca^2+^ leak. Thus, the concentration of Ca^2+^ in the cytosol is significantly decreased. The alteration of Ca^2+^ content induces a transient membrane depolarization and generates cell current for sweet and umami signaling transduction. RYR1 channel in bud tissue as a new one in regulation of sweet and umami signaling transduction is verified in this study. Additionally, a “metabolic clock” notion based on nutrient (sweet and umami) sensing mechanism was proposed.

**Table 1.**
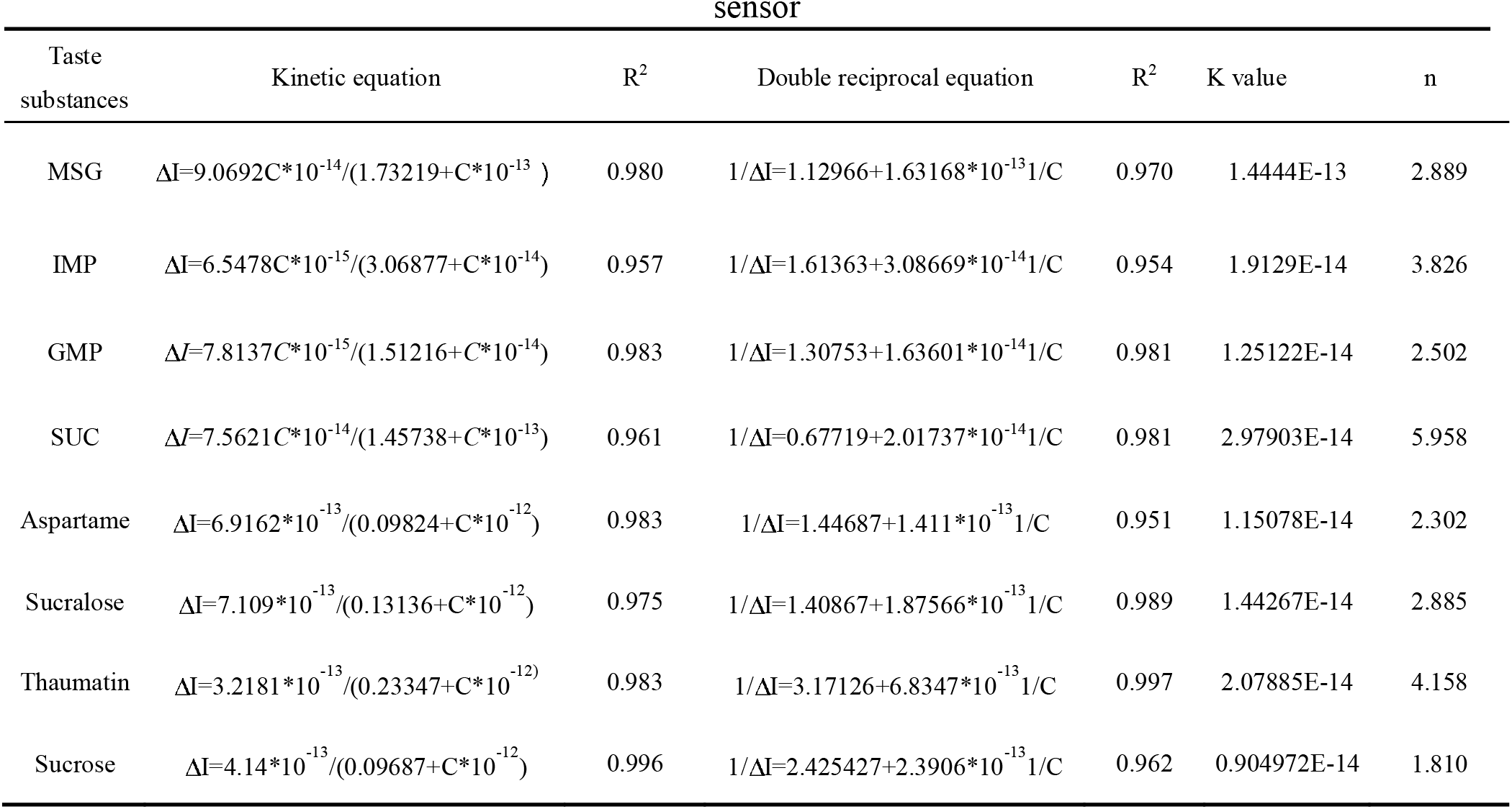
The K_a_ values of sweet and umami molecules based on rat taste bud tissue

## Acknowledgements

This work was supported by the National Natural Science Foundation of China (Grant No. 31901813, 31972198, 31671857, 31901782).

## Author contributions

Wenli Wang performed experiments, analysed data, wrote the paper. Dingqiang Lu performed experiments, analysed data. Qiuda Xu and Yulian Jin performed experiments. Guangchang Gang conceived the project, designed experiments, analysed data, provided funding. Yuan Liu conceived and supervised the project, provided funding.

## Appendix A. Supplementary data

**Supplementary data (Table S1)** to this article can be found online at

